# Extracellular riboflavin induces anaerobic biofilm formation in *Shewanella oneidensis*

**DOI:** 10.1101/2021.02.19.431833

**Authors:** Miriam Edel, Gunnar Sturm, Katrin Sturm-Richter, Michael Wagner, Julia Novion Ducassou, Yohann Couté, Harald Horn, Johannes Gescher

## Abstract

Some microorganisms can respire with extracellular electron acceptors using an extended electron transport chain to the cell surface. These organisms apply flavin molecules as cofactors to facilitate one-electron transfer catalysed by the terminal reductases and as endogenous electron shuttles. In the model organism *Shewanella oneidensis*, riboflavin production and excretion triggers a specific biofilm formation response that is initiated at a specific threshold concentration, similar to canonical quorum sensing molecules. Riboflavin-mediated messaging is based on the overexpression of the gene encoding the putrescin decarboxylase *speC* which leads to posttranscriptional overproduction of proteins involved in biofilm formation. Using a model of growth-dependent riboflavin production under batch and biofilm growth conditions, the number of cells necessary to produce the threshold concentration per time was deduced. Furthermore, our results indicate that specific retention of riboflavin in the biofilm matrix leads to localized concentrations which by far exceed the necessary threshold value.

**Importance:** Ferric iron is the fourth most abundant element of the earth crust. It occurs at neutral pH in the form insoluble iron minerals. The dissimilatory reduction of these minerals is an import part of global geological cycles and is catalyzed by microorganisms through extended respiratory chains to the cell surface. *Shewanella oneidensis* is one of the best understood model organisms for this kind of extracellular respiration. Flavins are important for the reduction of extracellular electron acceptors by *S. oneidensis*. since they have a function as (I) cofactors of the terminal reductases and (II) electron shuttles. In this study we reveal that flavin molecules are further employed as quorum sensing molecules. They are excreted by the organisms in a growth dependent manner and lead to anaerobic biofilm formation as a specific response at a certain threshold concentration. Although we know multiple examples of quorum sensing mechanisms, the use of riboflavin was so far not described and at least in *S. oneidensis* proceeds via a new regulatory routine that proceeds on the trancriptomic and posttranscriptomic level.

## Introduction

Flavin molecules accelerate extracellular respiratory processes. These processes are specific adaptations used by cells to couple oxidation of an electron donor to respiratory electron transport on insoluble or at least membrane-impermeable electron acceptors (1). These reduction processes are of environmental relevance. For instance, iron, the fourth most abundant element in the earth crust, occurs in soil and sediments in the form of insoluble iron oxides or oxyhydroxides and is one target for extracellular electron transfer processes. Moreover, reduction of insoluble electron acceptors can be applied in bio-electrochemical systems (BES), in which a solid-state anode is used by the microorganisms as an electron acceptor instead of environmental iron or manganese minerals. Hence, the organisms catalyse in BES the direct conversion of chemical into electrical energy.

Riboflavin and other flavin species are excreted by the γ-proteobacterium *Shewanella oneidensis*, which is a model organism for extracellular electron transfer (2). Extracellular flavins can be used as endogenous electron shuttle by *S. oneidensis*, and it was discovered that the presence of flavins has a positive impact on extracellular respiration kinetics (2–4). Recently, it was revealed that the influence of flavins is also due to their function as cofactors of the terminal reductases of the organism. As cofactor, riboflavin causes acceleration of the activity of outer membrane cytochromes by facilitating one electron transport via the formation of semiquinones (5–7). Meanwhile, it has become evident that flavins are not only cofactors of outer membrane cytochromes of *S. oneidensis* but also of *Geobacter sulfurreducens* (the other main model organism for extracellular electron transfer processes) (6, 8). Furthermore, it was demonstrated that flavo-enzymes are also used as terminal reductases of ferric iron in Gram-positive organisms (9).

Although the excretion of flavins was hypothesized to be of minor relevance regarding necessary energy and carbon source investment (2), it is still unclear why *Shewanella* strains do not bind flavin molecules as tightly to the outer membrane cytochromes as do *Geobacter* strains. A release of flavins into the medium supernatant has not been discovered for *Geobacter* strains so far. As a result of this study, we suggest that riboflavin in addition to its function as cofactor and potential electron shuttle is also a messenger or quorum sensing molecule used at least to facilitate anaerobic biofilm formation in *S. oneidensis*.

Quorum sensing is a microbial process that regulates the initiation of physiological responses in a cell concentration-dependent manner. The first process that was classified as quorum sensing-dependent was bioluminescence (10). In this process, it is particularly understandable that the energy-dependent process of light emission is initiated only when the density of organisms would allow the production of detectable light signals. Later, other processes were also revealed to be quorum sensing-dependent, including for instance, the production of pathogenicity factors or biofilm formation. Among Gram-negative organisms, four characteristics seem to be present in most quorum sensing systems. First, acyl-homoserine lactones or molecules synthesized from S-adenosylmethionine are mostly used as quorum sensing molecules. Second, the quorum sensing molecules can diffuse through the bacterial membranes. Third, quorum sensing changes the expression of a multitude of genes, and last but not least, quorum sensing leads to autoinduction of genes necessary for messenger molecule production (11). Nevertheless, exemptions from these rules have been presented many times. For instance, in *Shewanella baltica*, quorum sensing was discovered to be dependent on diketopiperazines, a class of small cyclic peptides (12). These compounds were previously identified as belonging to a family of molecules used for interspecies or interkingdom communication (13). Nevertheless, *S. baltica* relies on these substances for quorum sensing-induced pathogenesis while production of acyl-homoserines has not been detected so far.

In this study, we reveal that low riboflavin concentrations induce a specific transcriptomic response in a concentration-dependent manner in *S. oneidensis*. The addition of riboflavin triggers overexpression of one single gene, which encodes the ornithine-decarboxylase *speC*. The activity of the corresponding enzyme most probably leads to a proteomic response potentially based on putrescine-dependent regulation. Part of this response involves overproduction of proteins involved in biofilm production. Using data on riboflavin formation by planktonic and biofilm cells, we established a model that helped predict the necessary cell concentration and time to reach sufficient riboflavin concentration to start the respective physiological response of the organisms.

## Results

### Riboflavin addition triggers a specific transcriptomic response

Riboflavin addition causes enhanced biofilm and current production in BESs (14, 15). This process could occur for two different reasons: (1) addition of an electron shuttle enables biofilm cells to thrive more distantly localized to the anode or (II) external riboflavin triggers biofilm formation. In this case the higher cell density on the anode would, as a secondary effect, lead to higher current densities. In order to investigate this second hypothesis, a transcriptomic analysis was conducted using cells that were either grown with or without the addition of 37 nM riboflavin, a concentration that was also used in a previous study (14). To exclude the effects of riboflavin on gene expression due to its electron shuttling function and focus on its potential role as messenger molecule, fumarate, instead of an anode, served as electron acceptor.

Comparison of the two transcriptomes revealed that only one gene, *speC*, encoding for the ornithin decarboxylase of the organism was significantly upregulated with more than a 2-fold change (FDR p-value ≤ 0.05, Tab. 1). SpeC facilitates the production of the polyamine putrescine.

**Table 1:** Significantly regulated genes after riboflavin addition. Genes with a FDR p-value lower than 0.05 and a fold change higher than 2 or lower than −2 are shown. For all genes scored, the fold change was calculated by dividing the mutant value by the wild type value. If the number was less than one the negative reciprocal is listed.

| Gene | Fold change (FDR p-value)<br>riboflavin addition vs. no riboflavin addition |
| --- | --- |
| *speC* | 2.04 (1.92E-10) |
| *srtA* | -2.04 (4.24E-3) |
| *prpR* | -2.05 (6.61E-4) |
| *prpB* | -2.2 (2.86E-5) |

To prove whether an increase in expression of this gene causes higher currents and accelerated biofilm production after riboflavin addition in BES, the gene was expressed from an inducible plasmid in *S. oneidensis* wildtype. As depicted in figure 1, the positive effect of *speC*-overexpression on current production and biofilm formation was almost identical to the effect of adding 37 nM riboflavin. Moreover, *speC* deletion mutants were not capable of responding to the addition of this riboflavin concentration (Fig. 1). Furthermore, addition of 37 nM riboflavin to a strain that overexpresses *speC* did not lead to a further increase in cell number or current. Hence, at this concentration, potential electron shuttling by riboflavin cannot further accelerate electron transfer in cells that overexpress *speC*. Producing a similar effect of enhanced current production via the addition of putrescine to the medium was also attempted. Still, the addition of putrescine at a concentration range between 0.5 and 5 μM did not lead to a significant increase in current densities (Fig. S1).

**Fig. 1:**
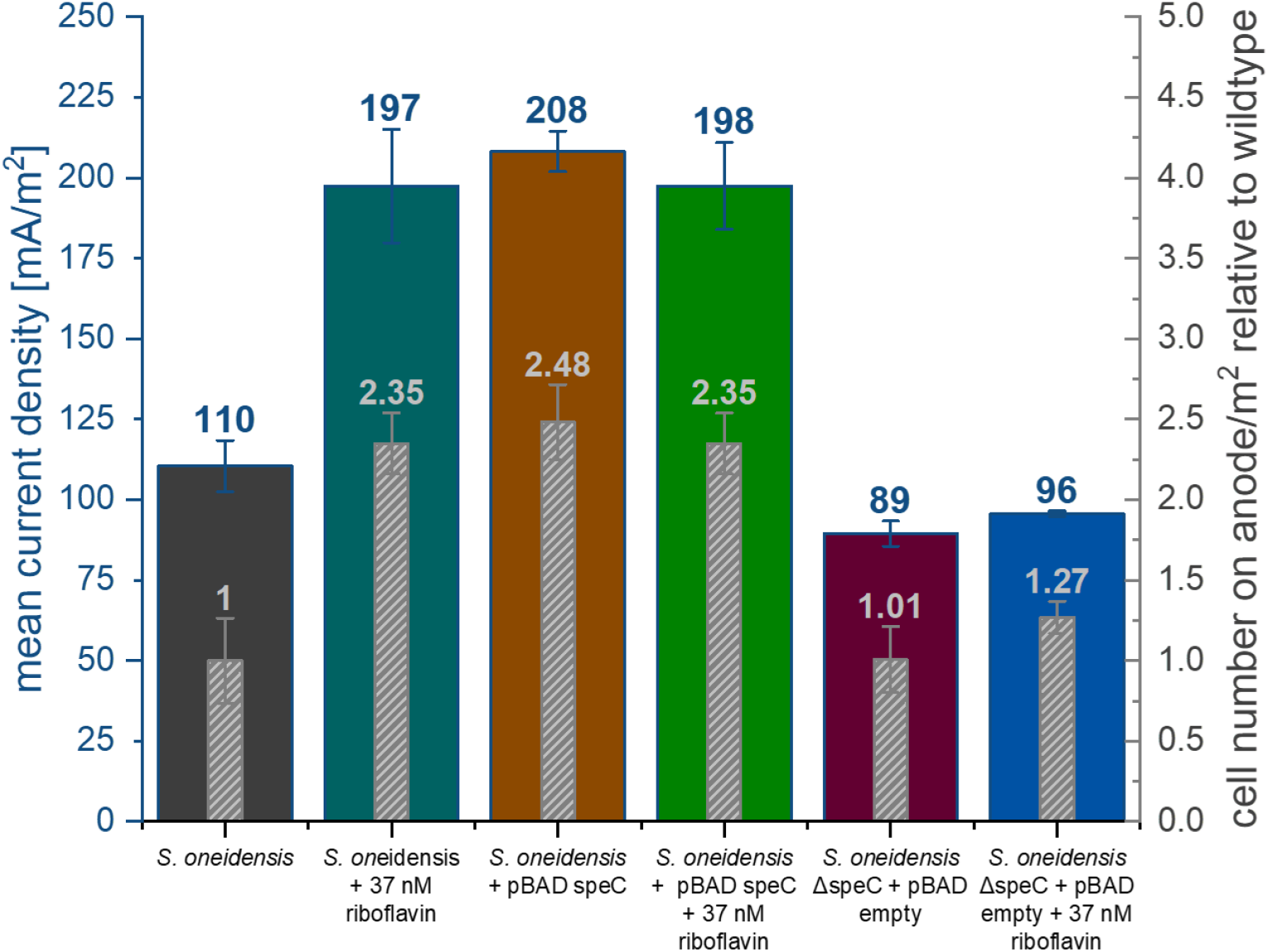
Impact of riboflavin and *speC* expression on current and biofilm formation on anode surfaces. Overexpression of *speC* resembles the phenotype after riboflavin addition, while deletion mutants seem to be blind for the riboflavin signal. The bar chart shows an increase in current density of 1.8-fold due to the addition of 37 nM riboflavin. Furthermore, the amount of cells on the anode increases 2.4-fold. A very similar effect can be observed by the overexpression of *speC*. The addition of 37 nM riboflavin to a *speC* overexpressing strain leads to no increase in current density or in cell number. The deletion of *speC* leads to a slight decrease in current density, but the supply of 37 nM riboflavin does not show any significant effect on the *speC* knockout strain. Error bars represent the standard deviation from individual replicates (n=3).

Besides *speC,* only three other genes were significantly regulated with a more than 2-fold change. The *srtA* gene was downregulated 2-fold. This gene encodes a putative sortase, a class of enzyme that catalyses the covalent attachment of specific proteins to the cell wall of Gram-positive and occasionally Gram-negative bacteria (16). Furthermore, the genes for *prpR* and *prpB* (2.1- and 2.2-fold, respectively), which are both involved in propionate degradation, were downregulated. Although the previous experiments revealed that *speC* overexpression was sufficient to completely mimic the effect of riboflavin addition, we also analyzed the potential effects of the sortase as its activity might have an effect on the surface chemistry of the organisms. Still, deletion of *srtA* gene from the genome of the organism did not lead to an increase in current density as would have been expected if the sortase was involved in the observed increased biofilm production (Fig. 2). Of note, it was so far not possible to generate deletion mutants in *prpR* and *prpB*.

**Fig. 2:**
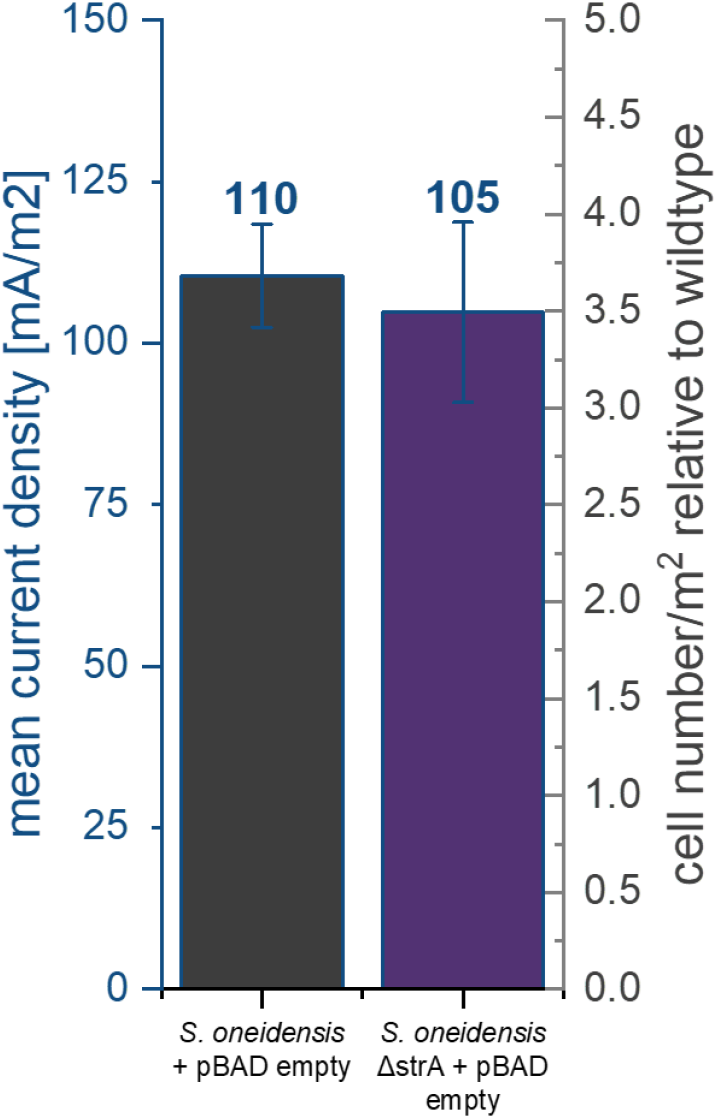
Impact of *srtA* deletion on current generation and biofilm formation on anode surfaces. The bar chart shows that the knockout of *srtA* does not have any significant effect on the current density. Error bars represent the standard deviation from individual replicates (n=3).

### Identifying the minimal riboflavin threshold for speC induction

Riboflavin is excreted by the cells during growth and its concentration consequently increases over time in a batch system (Fig. 4). Riboflavin concentrations in batch experiments that have been reported so far vary apparently depending on growth conditions between 30 and 450 nM (17–19). Since we hypothesized that riboflavin could be a quorum sensing molecule that is used by the cells to initiate a cell concentration-dependent physiological response, we quantified the response of the *speC* promoter on the addition of riboflavin at concentrations between 0 and 100 nM.

**Fig. 4:**
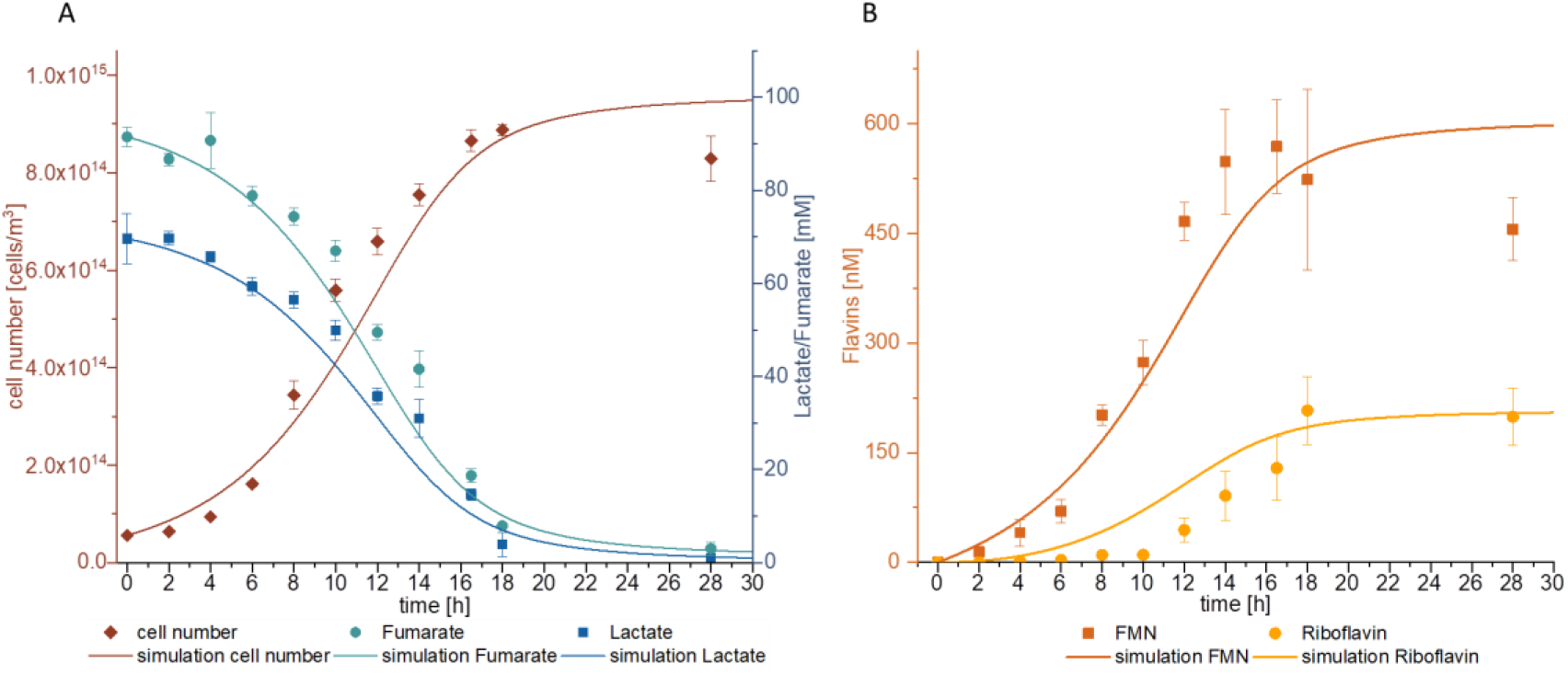
Growth (A) and flavin secretion (B) of *S. oneidensis* wildtype in a batch culture and results of the modelling attempt. Measured values are depicted as point squares or bars, while the modelling results are shown as solid lines. Error bars represent the standard deviation from 3 individual bacterial samples (n=3).

The expression of the *speC* gene was measured via quantitative polymerase chain reaction (PCR) as shown in Fig. 3. While the addition of 5-15 nM riboflavin showed a slight decrease in *speC*-expression, we observed a distinct upregulation after addition of more than 15 nM. A similar increase in *speC* expression was observed in a growth experiment without external addition of riboflavin. Expression increased as soon as the cellular riboflavin excretion led to medium concentrations of 15 nM (Fig. S3). Moreover, the response was specific to this flavin species as the addition of flavin-adenine-dinucleotide (FAD) or flavin-mononucleotide (FMN) could not trigger a similar response (Fig. 3).

**Fig. 3:**
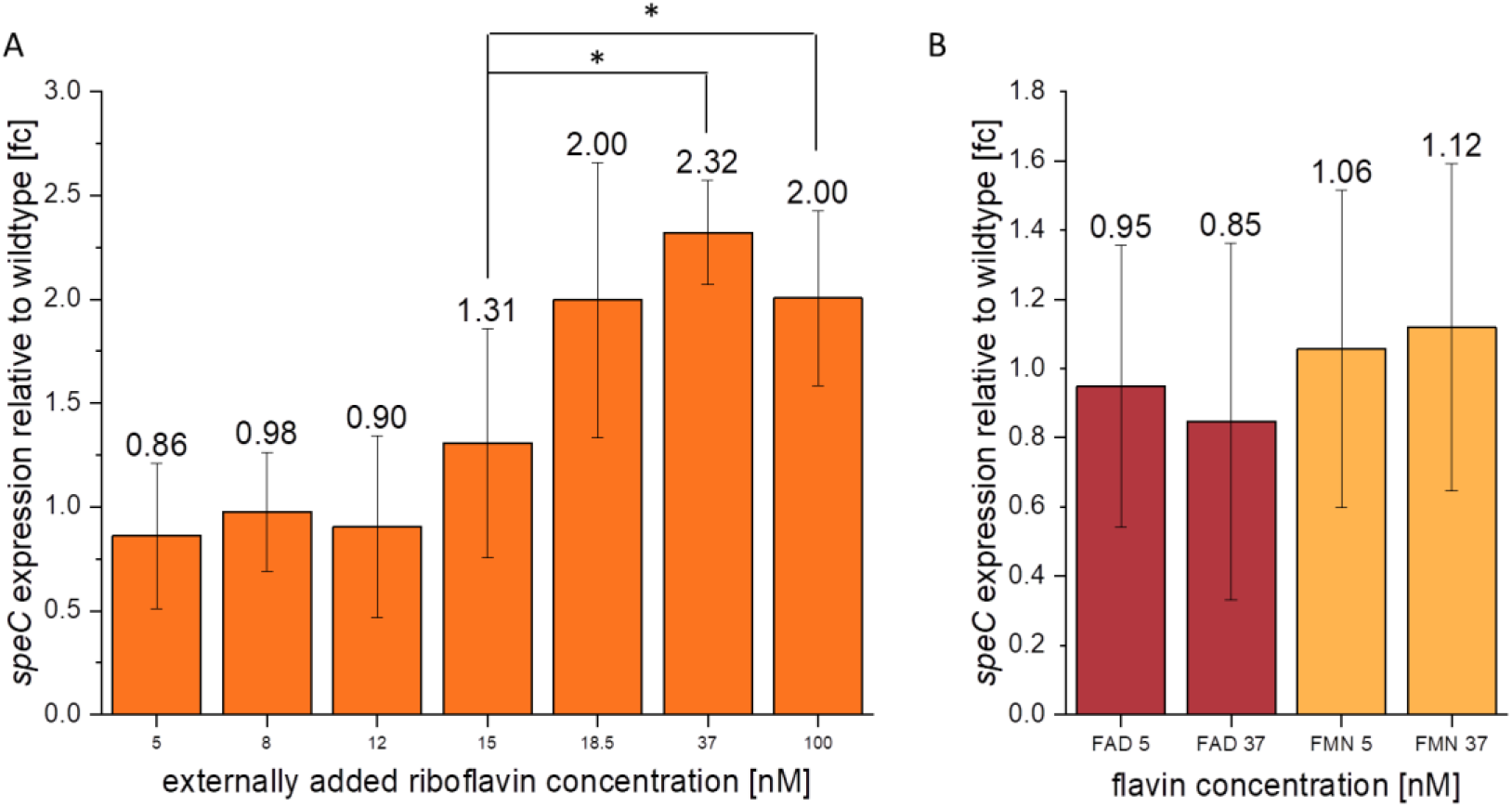
(A) *speC* expression after the addition of different concentrations of riboflavin relative to transcript abundance without riboflavin addition. The addition of up to 15 nM riboflavin does not have any significant effect on *speC* expression, while the addition of 18.5, 37 and 100 nM riboflavin leads to a 2- and 2.3-fold increase in *speC* expression, respectively. Error bars represent the standard deviation from individual replicates (n=3). (B) *speC* expression after the addition of different concentrations of flavin adenine dinucleotide (FAD) and flavin mononucleotide (FMN) relative to cells without exogenous flavin addition. The addition of FAD and FMN in these concentrations does not have a significant effect on *speC* expression. Expression is normalized to the gene for the RNA polymerase subunit *rpoA.* Error bars represent the standard deviation from individual replicates (n=3). Asterisks represent significant differences (unpaired *t*-test *p* < 0.05).

### Post-transcriptional effect of SpeC

So far, the experiments revealed that riboflavin triggers a concentration-dependent specific response of the *speC*-promoter, which in turn leads to biofilm production and enhanced current densities. Still, since no other gene was significantly upregulated in our study and since the genes that were downregulated also do not have a function regarding the observed biofilm phenotype, we asked whether the overexpression of *speC* might have an effect on the cell proteome. To this end, a quantitative proteomic study to compare the proteome of cells with and without *speC* overexpression was conducted. Again, the set of significantly differentially expressed proteins was limited (Tab. 2). Among them, we found that PuuA, which is involved in putrescine degradation, was more highly produced when *speC* was overexpressed. Conversely, less AguB, which is involved in putrescine production, was produced. Although many of the differentially produced proteins cannot be linked to the observed biofilm phenotype, we found at least six proteins that were previously reported to have a putative function that could explain increased biofilm formation. Two of the overproduced proteins, WbpP and ProQ, were previously shown to be important for biofilm or capsule formation (20–22). WbpP is a putative UDP-N-acetylglucosamine C4 epimerase. The corresponding gene seems to form an operon with *wbpA*, which encodes a UDP-D-GlcNAc dehydrogenase. WbpA was revealed to be slightly overproduced although with standard deviations that excluded statistical significance. Furthermore, ProQ, a posttranscriptional regulator in *E. coli* that positively affects biofilm formation (23), is overproduced. Among the proteins that were downregulated when *speC* was overexpressed, is SO_1208, a protein containing a GGDEF and an EAL-domain. Proteins containing these domains are often involved in adjusting intracellular levels of cyclic-di-GMP, a key messenger molecule for regulating biofilm formation. Furthermore, two putative proteases and a short chain dehydrogenase (SO_2766, SO_0491 and SO_1674) are under-expressed. Related proteins in other organisms were previously shown to be involved in cell detachment (24–26).

**Tab. 2:** Significantly regulated proteins due to *speC* expression. Proteins with a p-value lower than 0.05 and a log_2_ fold change greater than 1 or lower than −1 are shown.

| Protein | Log<sub>2</sub> fold change (p-value)<br>*speC* expression vs wildtype |
| --- | --- |
| SpeC | 9.65 (5E-06) |
| SO_3964 | 4.09 (0.0445) |
| ArgB | 3.22 (2E-06) |
| SO_0576 | 2.5 (2E-05) |
| DctP | 2.32 (0.0026) |
| IvdG | 2.32 (0.0208) |
| ProQ | 2.08 (0.0308) |
| SO_3698 | 1.75 (0.0426) |
| ThiL | 1.7 (8E-06) |
| SO_4371 | 1.67 (0.0215) |
| GarR | 1.64 (0.0005) |
| PuuA | 1.53 (0.0002) |
| AdhB | 1.53 (0.0003) |
| LeuB | 1.49 (0.0117) |
| Tpx | 1.46 (0.0175) |
| HemL | 1.4 (0.0003) |
| YebR | 1.31 (0.0445) |
| MetH | 1.26 (0.0349) |
| WbpP | 1.23 (0.0004) |
| SelD | 1.14 (0.0009) |
| RmlA | 1.09 (0.0017) |
| Upp | 1.06 (0.0045) |
| TrpA | 1.03 (9E-05) |
| SO_2766 | -1.09 (0.0047) |
| SO_1059 | -1.27 (0.0114) |
| AguB | -1.29 (0.0008) |
| SO_4378 | -1.33 (0.0004) |
| SO_A0051 | -1.35 (0.0004) |
| SO_0695 | -1.43 (0.0262) |
| Rnr | -1.46 (0.0007) |
| SO_4429 | -1.57 (0.0309) |
| SO_0491 | -1.66 (0.0195) |
| SO_3119 | -1.97 (6E-05) |
| SO_2636 | -2.01 (0.0027) |
| SO_1208 | -2.02 (0.0021) |
| SO_1674 | -2.25 (0.0218) |
| RpoE | -2.69 (0.0391) |

### Modelling riboflavin production in a batch culture

Our next aim was to assess the necessary cell concentrations and times to reach the riboflavin threshold concentration for biofilm production (Fig. 4). Hence, the kinetic and stoichiometric values for riboflavin production were determined from a culture growing anoxically with lactate and fumarate (Fig. 4). The data depicted in figure 4 corroborates data from other research groups demonstrating that FMN is released by the cells and gets slowly converted into riboflavin. The latter reaction is supposed to proceed abiotically. Nevertheless, a comparison of the FMN and riboflavin concentrations after 18 and 28 h suggests that active growth is required for this reaction as the riboflavin concentration does not significantly increase within this time frame although sufficient amounts of FMN are still available.

Using the data from figure 4, we formulated a model for the production process that was applied to determine a time/cell number dependency for the accumulation of 18.5 nM riboflavin (see Tab. S3). The model (supplementary material) describes cell growth based on Monod terms with an electron acceptor (fumarate) and electron donor (lactate). FMN production is described as a growth-related process, whereas the formation of riboflavin from FMN occurs according to first-order kinetics. We had to combine the first-order formation with the two above mentioned Monod terms as the experimental results indicate riboflavin production stops as soon as the substrate is depleted.

Using this model, it was possible to predict the time required for a culture of a certain density to produce the necessary 18.5 nM riboflavin to trigger the biofilm formation response (Fig. 5). The estimated time intervals seem reasonable if the cells could grow in an environmental setting without or with slow mixing.

**Fig. 5:**
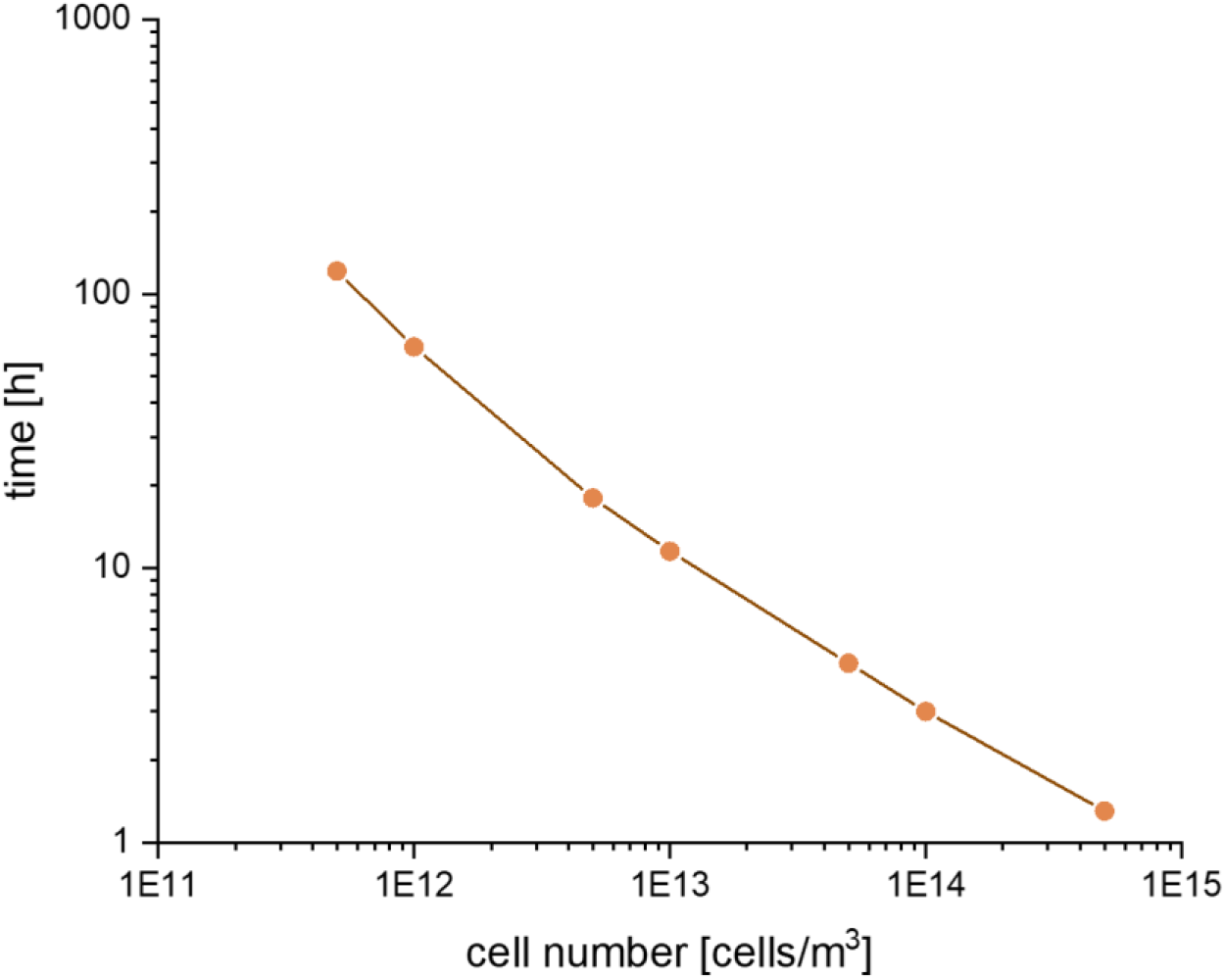
Exemplary prediction of ime needed to produce 18 nM riboflavin with different cell numbers. The delineation is based on the model described in the text and supplemental material.

### Realization of riboflavin-based quorum sensing is possible in a biofilm setup

Planktonic growth in open water and not in a laboratory batch system will always cause dilution of the excreted riboflavin to values below the cellular detection limit. Consequently, the dependencies described above will be applicable in nature only under conditions in which an enclosed environment develops, for instance, in a lagoon setting. Still, the natural way of living of most microorganisms is in the form of a biofilm. Hence, we asked whether retention of riboflavin in a *S. oneidensis* biofilm or more specifically in its extracellular polymeric substances (EPS) could be sufficient to reach necessary riboflavin concentrations.

The *S. oneidensis* biofilm was grown in a microfluidic device under constant flow. Finally, a mean biofilm thickness of 76.3 ± 12.9 μm was reached (Fig. 6). After 121 h, no measurable lactate in the effluent was found. In contrary to the batch experiment, after 97 h riboflavin was the dominant flavin species in the effluent of the system. After 143 h, the biofilm was lysed, and the flavin content was measured. Using optical coherence tomography (OCT)-data for biofilm volume, it was possible to determine the concentration of the different flavin species. The biofilm contained 3670 ± 224 nM riboflavin, 213 ± 19 nM FMN, and 292 ± 47 nM FAD. Hence, the observed flavin concentration was far above the identified concentration for riboflavin-based biofilm production. We sought to determine only the extracellular riboflavin concentration by carefully resuspending the developed biofilm. Nevertheless, we also measured the intracellular riboflavin concentration to exclude the possibility that partially lysed cells could have greatly contributed to the riboflavin concentration. The riboflavin content within the cells was at a level of 2.81 × 10^−3^ fM per cell. Thus, 5.15 × 10^10^ cells/ml (as in the analysed biofilm) would contain 144.72 nM of intracellular riboflavin, which is more than one magnitude below the measured value of 3670 nM. This finding agrees with the observation of von Canstein and colleagues, who could only measure trace amounts of riboflavin in the cytoplasm (27).

**Fig. 6:**
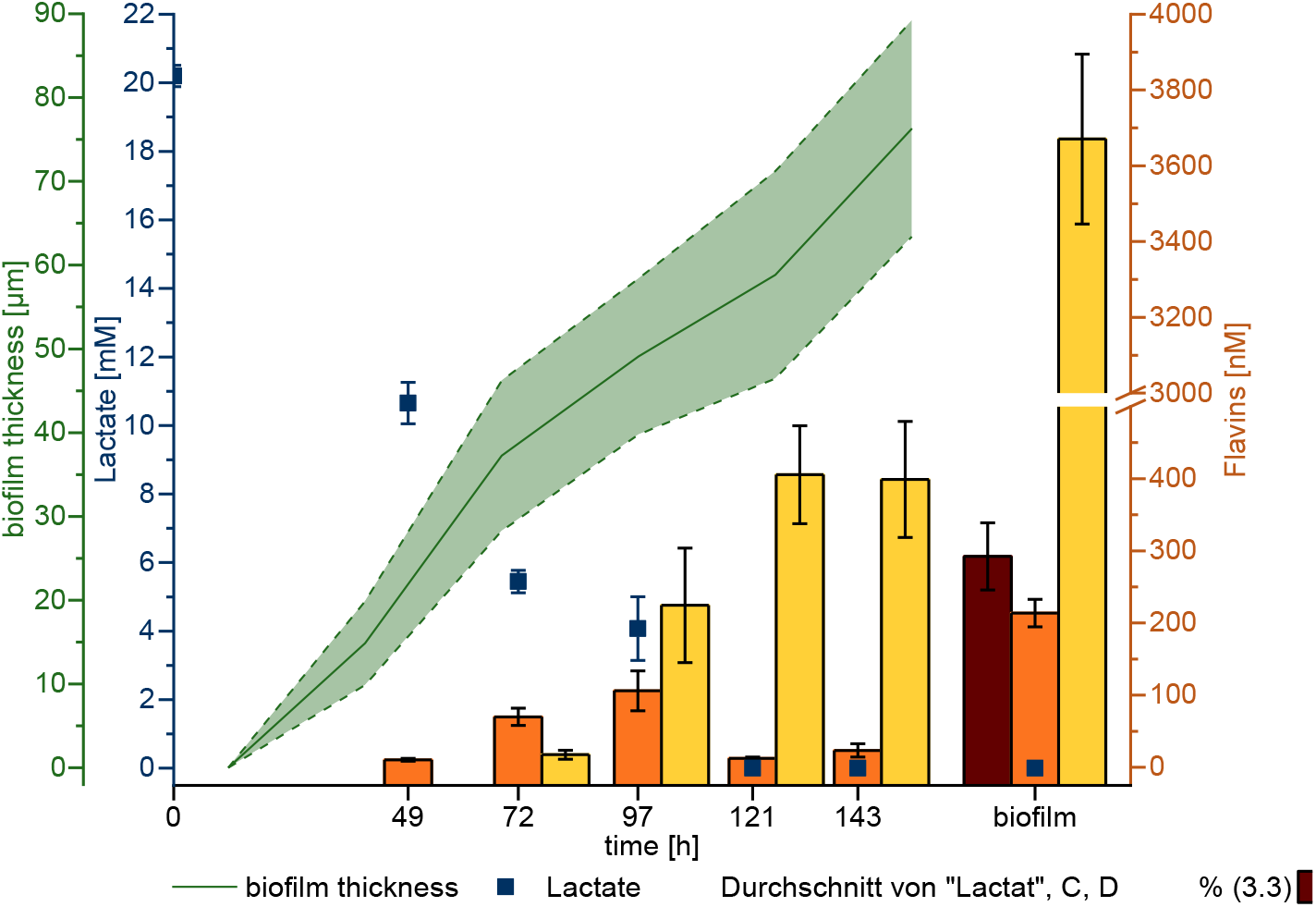
Mean biofilm thickness, lactate and flavin concentration of *S. oneidensis* wildtype grown in a PDMS chip over time. After 121 h there was no detectable amount of lactate in the effluent. The flavin content significantly increased over time. However, especially the riboflavin concentration is much higher within the biofilm compared to the effluent. Error bars represent the standard deviation from 3 individual bacterial samples (n=3).

We also aimed to model riboflavin production in the biofilm system based on the quantitative data depicted in figure 6. Model description and simulation results are provided in the supplementary material. For our purpose, the biofilm model was mainly used as analytical tool. In fact, the simulation results (Fig. S4) show one clear result. The release of product (riboflavin) from biofilm into the bulk medium cannot solely be explained by a traditional reaction diffusion approach for biofilm systems (28). Obviously, part of riboflavin product is not directly released into the bulk liquid. To fit the simulation with the achieved results, it was necessary to reduce the diffusion coefficient of riboflavin in water by three orders of magnitude. Moreover, yield coefficients for biomass and product and the first-order reaction constant for FMN transformation to riboflavin had to be adapted and compared to the values measured in the batch culture (see Tab. S3). We assumed that the two orders of magnitude higher cell density within the biofilm had an impact on yield coefficient for FMN and first-order reaction rate for riboflavin from FMN.

Biofilm thickness and lactate consumption in the flow cell can be displayed very satisfactorily with the chosen model. However, for both FMN and riboflavin, the simulation shows release from the biofilm into the bulk liquid starting already on the first day. The experimental results clearly show that the cells do not release the products that fast. The decreased diffusion coefficient for riboflavin allows for an adequate simulation of its concentration within the biofilm after 140 h (Fig. S4 e). However, the concentrations of riboflavin in bulk and biofilms can be misleading. In fact, the total amount of riboflavin within the biofilm after 140 h is only 4.6 × 10^−3^ nmol, and the riboflavin produced by the biofilm (and transported out of the system) totals 24 nmol.

## Discussion

This study revealed that *S. oneidensis* cells can sense the extracellular riboflavin concentration and use this information to initiate an intracellular process leading to biofilm formation. This process is apparently based on a posttranslational overproduction of proteins triggered by the intracellular activity of SpeC. The post-translational response involves WbpP, a key protein for extracellular matrix production (20, 21). Modelling and biofilm experiments showed that the necessary riboflavin concentration in biofilms can be reached with a low number of attached cells. Moreover, comparing the model with experimental data from the biofilm analysis clearly indicates a specific retention of riboflavin within the biofilm matrix.

The observed riboflavin-dependent response of the cells is not in line with canonical quorum sensing systems. It does not operate with an acylhomoserine lactone, and autoinduction of the signalling cascade could not be observed. Moreover, the final response seems to be triggered post-translationally and does not directly depend on the transcription level. Nevertheless, riboflavin is produced in a growth-dependent manner and leads after a distinct threshspecifically recognized by the cells. Moreover, the physiological biofilm formation response is similar to other quorum sensing-based systems. Hence, we hypothesize that riboflavin should be added to the list of quorum sensing molecules. As mentioned above, other non-canonical quorum sensing mechanisms have already been described in other *Shewanella* species. Thus, the use of alternative quorum sensing mechanisms might be a general characteristic of the genus. Still, a link between riboflavin as potential quorum sensing molecule and the canonical N-acyl homoserine lactone messengers was provided in a study by Rajamani *et al*. (29). The authors observed that cultures of the alga *Chlamydomonas* produced and secreted lumichrome into the culture supernatant. Lumichrome is a derivative of riboflavin. Moreover, the authors detected that lumichrome and riboflavin could stimulate the quorum sensing receptor LasR from *Pseudomonas aeruginosa*. Hence, riboflavin could be a ubiquitous communication molecule that operates among different kingdoms (29).

Ding and colleagues observed the impact of polyamines on aerobic biofilm production by *S. oneidensis* (30). Using a transposon screen, the authors discovered a mutant that apparently produced more EPS compared to the wild type. Detailed analysis revealed a 2-fold increase in EPS production and that the gene encoding the ornithin decarboxylase *speF* (an isoenzyme of SpeC) was interrupted by transposon integration. The mutant was hyper-adherent and formed tower-like biofilms when examined in static biofilm assays. Still, Ding and colleagues conducted experiments under oxic conditions, and the requirements for an efficient anaerobic biofilm system thriving with an insoluble electron acceptor seem to be quite different. For instance, EPS production *per se* can have also an insulating effect as shown in a variety of studies (31–33). Moreover, efficient electron transfer demands confluent coverage of the biofilm substrate and not tower-like biofilm structures as observed by Ding et al. (30). The latter will simply not lead to an optimized distance between the bulk of the cells and the electron acceptor. Nevertheless, we tested also the influence of *speF* on current production in a BES. As expected, a marker-less *speF* mutant produced less current compared to the wild type (Fig. S2).

The observed cellular response was riboflavin-specific as other flavin molecules did not lead to *speC*-overexpression. Flavin excretion starts with the export of FAD from the cytoplasm by the specific transporter Bfe. In the periplasm, FAD is used as cofactor and to a certain extend hydrolyzed to FMN and AMP by the periplasmic 5’ nuleotidase UshA (34). In line with the UshA activity in the periplasm, FMN is the main flavin species under planktonic conditions followed by significantly lower levels of riboflavin (3, 27, 34). Hence, the cells do not detect the flavin molecule with the highest concentration but the molecule that is formed by slow hydrolysis of FMN. This process causes a time delayed cellular response to flavins in the environment, which might have evolved to avoid a cellular response that is too rapid. In fact, evolution might have selected a response that is dependent on a diffusion-limited or batch environment in which the conditions are stable enough to allow enough time for FMN-hydrolysis to proceed and sufficient concentrations of riboflavin to accumulate. In a biofilm setting, the cells apparently evolved further measures to increase the local riboflavin concentration by retention of this substance. Therefore, already low cell numbers will trigger the SpeC-dependent biofilm formation response.

The results favor a post-transcriptional impact of putrescine on protein production. Of note, this kind of regulatory function of putrescine has so far only been observed for the gene cluster involved in putrescine degradation in *E. coli*, while a rather global impact on gene regulation was observed only in eukaryotic organisms (35–37). To date, it can only be assumed that putrescine binding to proteins or RNA will affect translation or the stability of proteins and mRNA involved in the observed response. Of great importance for the biofilm formation response might be WbpP, which was shown to be necessary for capsule formation in different strains (20, 21). The biochemical pathway for capsule formation has been widely studied in *Pseudomonas* strains. Here, glucose-1-phosphate is converted to UDP-*N*-acetyl-D-glucosamine, which is the central precursor of surface-associated carbohydrate synthesis. UDP-*N*-acetyl-D-galactosamine is then formed by the WbpP-catalyzed C4 epimerization of UDP-*N*-acetyl-D-glucosamine (38, 39). Dehydrogenation of the latter leads to the production of UDP-*N*-acetyl-D-galactosaminuronic acid. WbpP together with WbpA seems to form an operon in *S. oneidensis*. The latter converts UDP-*N*-acetyl-D-glucosamine to the corresponding uronic acids (22, 40). Hence, both enzymes compete for the same substrate. The observed post-transcriptional regulation leading to a higher production of WbpP compared to WbpA might be a way to fine tune the chemical composition of the capsule polysaccharide. Within the set of over- and underproduced enzymes, PuuA and AguB, respectively, were also detected. Hence, the cell also initiates countermeasures for putrescine production using SpeC-based regulation, which would likely be necessary for cellular homeostasis of this polyamine.

We developed a model for riboflavin production with the aim of specifying the number of organisms and the volume in which the organisms have to thrive to accumulate the necessary amount of riboflavin. The quantification of flavins in the biofilm-matrix indicate a rather high concentration that could not be correlated with the results from the planktonic experiments and the model based on the riboflavin diffusion coefficient, the average biofilm volume, and the average cell number per biofilm volume. The most probable explanation for this experimental result is that riboflavin is bound by the cells and/or the biofilm matrix. As cofactors, flavins are attached to the outer membrane cytochromes on the cell surface. Previous research revealed that the number of outer membrane cytochromes on the cell surface is between 1000 and 30,000 proteins per μm^2^ (41, 42). Using the average dimension of *Shewanella* cells (6.79 μm^2^) revealed by Sturm *et al*. (43) and the average number of cells per mL biofilm (5.15 × 10^10^), we can assume that the biofilm will contain between 3.49 × 10^14^ and 1.05 × 10^16^ outer membrane cytochromes per mL. This number is within the range of the measured riboflavin concentration of 3670 ± 224 nM riboflavin; which would be equivalent to 2.21×10^15^ riboflavin molecules per mL. Hence, outer membrane cytochrome-based retention of riboflavin likely allows for riboflavin-based messaging even under environmental conditions, such as those characterized by constant mixing. In other words, the observed process probably allows quorum sensing under conditions that would not lead to a cellular response with a canonical quorum sensing molecule that is quickly washed out of the biofilm by the medium flow.

## Materials and Methods

### Media and growth conditions

All strains used in this study are listed in table S1. *S. oneidensis* was precultured under oxic conditions in M4 minimal medium (pH 7.4) with 12 mM HEPES buffer and 70 mM lactate as electron donor. The medium was prepared as previously described (14). After 8 h of oxic growth, *S. oneidensis* was transferred to anoxic M4 medium containing 70 mM lactate as electron donor and 100 mM fumarate as electron acceptor. Oxygen was removed from the media via the repeated application of a vacuum and following sparging of the headspace with N_2_. The initial OD_600_ of the anoxic culture was set to 0.05. For genetic modifications, the strains were grown in LB medium under oxic conditions at 37 °C (*E. coli)* or 30 °C (*S. oneidensis*). If necessary, 2,6-diaminopimelic acid (DAP; 0.3 mM), kanamycin (Km; 50 μg/mL), and arabinose (0.1 mM) were added to the medium.

### Construction of expression plasmids

Plasmid pBAD202 was used for overexpression of the gene *speC*. The plasmid was cleaved using NcoI and PmeI. *speC* was amplified from the *S. oneidensis* wildtype genome. By using elongated primers (Primers 1 and 2, Tab. S2), an overlap to the plasmid was added that was used for isothermal ligation as described by Gibson and colleagues (44). The respective plasmid was verified by PCR analysis (Primers 3 and 4; Tab. S2) and subsequent sequencing.

### Construction of markerless deletion mutants

Marker-less deletion of genes was conducted according to Schuetz *et al*. (45). The respective pMQ150-based suicide vectors were also gained using isothermal *in vitro* recombination (44). The pMQ150 vector was cleaved using BamHI and SalI. Fragments with lengths of 500 bp up- and down-stream of the respective genes were amplified using primers 5–8 for *speC* deletion, primers 9–12 for *srtA* deletion, primers 13-16 for *prpR* deletion and primers 17-20 for *prpB* deletion (Tab. S2). The fragments contained an overlap to the pMQ150 plasmid in addition to each other. The three fragments were used for isothermal ligation as described by Gibson and colleagues (44) and transformed into the *E. coli* mating strain WM3064 (kind gift of W. Metcalf, University of Illinois). The suicide vector was then conjugated into *S. oneidensis*. All further steps were conducted as described by Saltikov and Newman (46). The respective deletions were verified by PCR analysis (Primers 21 to 26; Tab. S2) and subsequent sequencing. For the markerless deletion of prpR and prpB only the integration of the pMQ150-based vectors in the *S. oneidensis* genome was successful. The deletion of the residual genes could not be accomplished.

### Bioelectrochemical system (BES)

All BES experiments were conducted in triplicate using a single chamber BES with a working volume of 270 ml (33). Graphite felt (projected area of 36 cm^2^, SGL Group, Germany) and platinum mesh (projected area of 1.25 cm^2^, chemPUR, Germany) were used as working and counter electrode materials, respectively, and an Ag/AgCl electrode (Sensortechnik Meinsberg, Germany) served as the reference electrode. Prior to use, the working electrode was rinsed with isopropanol followed by deionized water. The complete BES setup was sterilized by autoclaving.

Before inoculating the reactors with microorganisms, cells were harvested from the anoxic preculture by centrifugation (7 min, 6000 g) and washed three times with medium not containing electron donor or electron acceptor. Afterwards, the cells were resuspended to a final OD_600_ of 0.07 in M4 medium containing 70 mM lactate. For pBAD-plasmid containing strains 50 μg/l kanamycin and 0.1 mM arabinose were added.

During chronoamperometric experiments, the working electrode was poised to 0 mV versus a standard hydrogen electrode (SHE), and current was monitored for 46 h. BES setups were constantly flushed with N_2_ gas in order to provide anoxic conditions. The medium was stirred constantly for thorough mixing.

### Flavin quantification

For quantification of flavins, liquid samples from anoxic *S. oneidensis* wildtype cultures were taken at different time points. The samples were filtered through a 0.2 μm-pore-size filter and analysed by reversed-phase high-performance liquid chromatography ([HPLC]; Luna® 5 μm C18(2) 100 Å, 250 × 4.6 mm column). The compounds were separated as previously described by van Canstein and colleagues (19) using a flow rate of 0.5 ml/min. The concentration of flavins was determined using a fluorescence detector at 440 and 525 nm excitation and emission, respectively.

### DNA Isolation for qPCR

The innuPREP Stool DNA Kit (Analytic Jena, Germany) was used to isolate DNA according to the manufacturers’ suggestions with minor modifications. Anodes from the BES setups were sliced into pieces, and 5 ml of SLS buffer (Analytic Jena, Germany) was added. The samples were vortexed vigorously for 1 min. Thereafter, the samples were incubated at 95 °C for 15 min. The samples were vortexed every 5 min. Afterwards, DNA isolation proceeded according to the manufacturer’s protocol. Relative cell quantifications of anode samples were based on three individual BESs. Two slices of one anode were used as a technical duplicate. Hence, every quantitative PCR result is based on the quantification of six samples.

### qPCR

For quantitative analysis of cell adhesion to the anode, a standard curve using biological triplicates of *S. oneidensis* wildtype in six different dilutions was established. Before isolation of the DNA, the cells were counted in two different dilutions (Neubauer chamber improved, Friedrichsdorf, Germany). Quantitative PCR (qPCR) was conducted using primers 27 and 28 (Tab. S2) according to Dolch *et al.* (47). For quantification of *speC* transcripts, *S. oneidensis* wildtype was grown in anoxic M4 minimal medium with the addition of 0, 5, 12, 15, 18.5, 37 and 100 nM riboflavin. The cells were harvested after 5 h of growth. Furthermore, cells were grown in anoxic M4 media without the addition of riboflavin and samples were taken after 30, 60, 180, 300, 480 and 600 min of growth. From all samples the RNA was isolated using the RNeasy Mini Kit (Qiagen, Germany) according to the manufacturer’s instructions. In order to remove any residual DNA, the samples were treated with RNase free DNase (Qiagen, Germany) overnight. Afterwards, the RNA was transcribed into cDNA using the iScript™ Select cDNA Synthesis Kit with random primers (Biorad, Germany). The amount of *speC* transcripts was quantified using primers 29 and 30 (Tab. S2) in a qPCR. To normalize the amount of *speC* transcripts, transcripts of the housekeeping gene *rpoA* were quantified (primers 31 and 32, Tab. S2).

### Transcriptomic analysis

Transcriptomic analyses were conducted with cells that were inoculated in anoxic M4 minimal media with 70 mM lactate as electron donor and 100 mM fumarate as electron acceptor. The starting OD_600_ was 0.2. After 5 h of growth, cells were harvested by centrifugation (7 min, 6000 g, 4°C). The supernatant was discarded, and pellets were frozen using liquid N_2_. For each analysis, two biological triplicates were combined into one sample, resulting in two transcriptomic samples per condition.

mRNA extraction and Illumina sequencing was conducted by IMGM Laboratories (Martinsried, Germany). Bioinformatic analysis was performed using the CLC Genomics Workbench version 12.0.3 (https://www.qiagenbioinformatics). Prior to mapping, low-quality reads were trimmed. The 75 nt single end sequencing reaction yielded 15 million reads on average (after trimming) mapping concordantly to the *S. oneidensis* MR-1 reference genome. The coverage was 233-fold. Transcript per kilobase million (TPM) values were used for comparison, and false discovery rates (FDR) p-values ≤ 0.05 account for statistical significance.

### Mass spectrometry-based quantitative proteomic analysis

Proteomic analyses were conducted with cells that were inoculated in anoxic M4 minimal media with 70 mM lactate as electron donor and 100 mM fumarate as electron acceptor. The starting OD_600_ was 0.2. After 5 h of growth, cells were harvested by centrifugation (7 min, 6000 g, 4°C). The supernatant was discarded, and pellets were resuspended in TRIS buffer (pH 6.8). Cells were lysed by two passages through a French Press.

The protein content of samples was quantified using the Bradford assay (48) with bovine serum albumin as standard. The protein concentration was then adjusted to 1 μg/μl.

Cell lysates were mixed with Laemmli buffer (49) and then stacked in a single band in the top of a sodium dodecyl sulfate-polyacrylamide gel electrophoresis (SDS-PAGE) gel (4%–12% NuPAGE gel, Invitrogen, Illkirch-Grafenstaden/France). After staining with R-250 Coomassie Blue (Biorad, Schiltigheim/France), proteins were digested ingel using modified trypsin (sequencing grade, Promega, Charbonnières les Bains/France) as previously described (50).

Resulting peptides were analyzed by online nano-liquid chromatography coupled to tandem mass spectrometry (MS) (Ultimate 3000 RSLCnano and Q-Exactive HF, Thermo Scientific, Illkirch-Grafenstaden/France). For this purpose, peptides were sampled on a 300 μm × 5 mm PepMap C18 precolumn (Thermo Scientific, Illkirch-Grafenstaden/France) and separated on a 75 μm × 250 mm C18 column (Reprosil-Pur 120 C18-AQ, 1.9 μm, Dr. Maisch, Ammerbuch-Entringen) using a 180-min gradient. The MS and MS/MS data were collected by Xcalibur (version 4.1.31.9, Thermo Scientific, Illkirch-Grafenstaden/France). Peptides and proteins were identified by Mascot (version 2.6.0, Matrix Science, London/United Kingdom) through concomitant searches against the Uniprot database (*S. oneidensis* MR-1 taxonomy, January 2019 version), a homemade classical contaminant database, and the corresponding reversed databases. Trypsin/P was chosen as the enzyme, and two missed cleavages were allowed. Precursor and fragment mass error tolerances were set at respectively at 10 and 25 mmu. Peptide modifications allowed during the search were carbamidomethyl (C, fixed), Acetyl (Protein N-term, variable) and oxidation (M, variable). The Proline software (51) was used to filter the results with conservation of rank 1 peptides, peptide score ≥ 25, peptide length ≥ 7 amino acids, FDR of peptide-spectrum-match identifications < 1% as calculated on peptide-spectrum-match scores by employing the reverse database strategy, and a minimum of 1 specific peptide per identified protein group. Proline was then used to perform a compilation, grouping and MS1 quantification of the validated protein groups.

The statistical evaluation was performed using the ProStar software (version 1.14) (52). Proteins identified in the reverse and contaminant databases, and proteins quantified in less than three replicates of one condition were discarded from the list.

After log2 transformation, abundance values were normalized by the vsn method before missing value imputation (slsa algorithm for partially observed values in the condition and DetQuantile algorithm for totally absent values in the condition). The statistical testing was conducted using *limma*. Differentially expressed proteins were sorted out using a log2 (fold change) cut-off of 1 and a p-value cut-off of 0.05, allowing an FDR inferior to 5% according to the adjusted Benjamini-Hochberg (abh) estimator to be obtained.

### Quantification of the biofilm volume via optical coherence tomography (OCT)

To quantify the average concentration of cells and riboflavin within a biofilm or the biofilm-matrix, respectively, *S. oneidensis* cells were grown in M4 minimal medium under oxic conditions using 20 mM lactate as electron donor. The cells were cultured in triplicate in a microfluidic flow-through system with straight polydimethylsiloxane (PDMS) channels and a total volume of 354 μL (53). The microfluidic systems were inoculated for 2 h using a culture with an OD_600_ of 0.08 and a flow rate of 3 ml/h. During biofilm growth, the pump rate was decreased to a rate of 1 ml/h. Biofilm growth was determined using OCT. After six days of growth, the supernatant of the biofilm was carefully removed, and the biofilm was resuspended by pipetting. Resuspended cells were counted (Neubauer chamber improved), and flavin content of the resuspended biofilm was analysed via HPLC. Furthermore, the volume of the total biofilm was quantified using a GANYMEDE spectral domain OCT system (Thorlabs GmbH, Dachau, Germany). Biofilm volume was determined at the beginning, the middle and the end of the three PDMS chips and the mean biofilm volume was calculated.

## Acknowledgments

This work was supported by a grant of the Bundesministerium für Bildung und Forschung (BMBF), no. 031B0847A Proteomic experiments were partly supported by the ProFI grant (ANR-10-INBS-08-01).

## Notes

### Competing Interest Statement

The authors have declared no competing interest.

