## Supplementary figures and images for "Extracellular riboflavin induces anaerobic biofilm formation in *Shewanella oneidensis*"

### Supplemental Figure 1

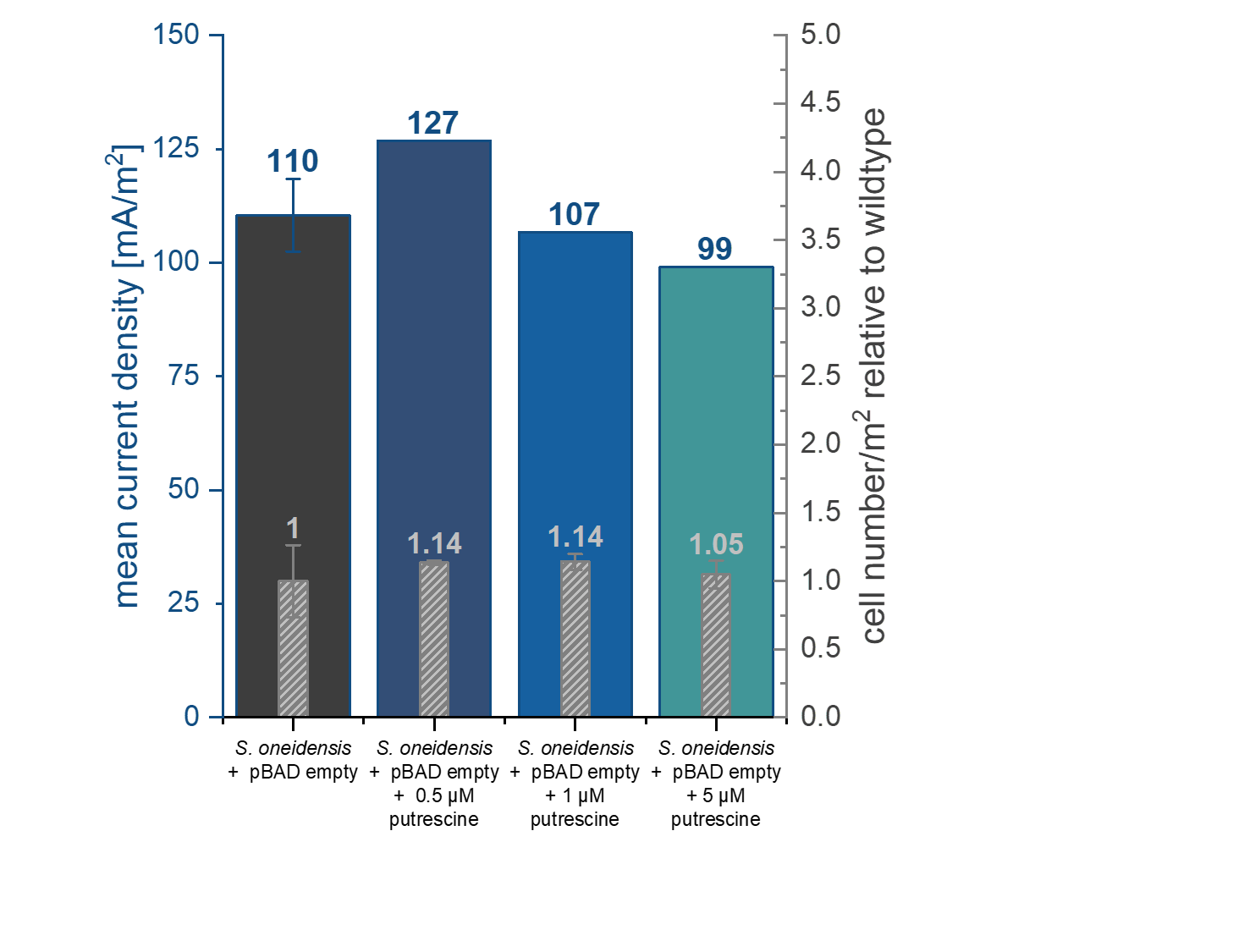

### Supplemental Figure 2

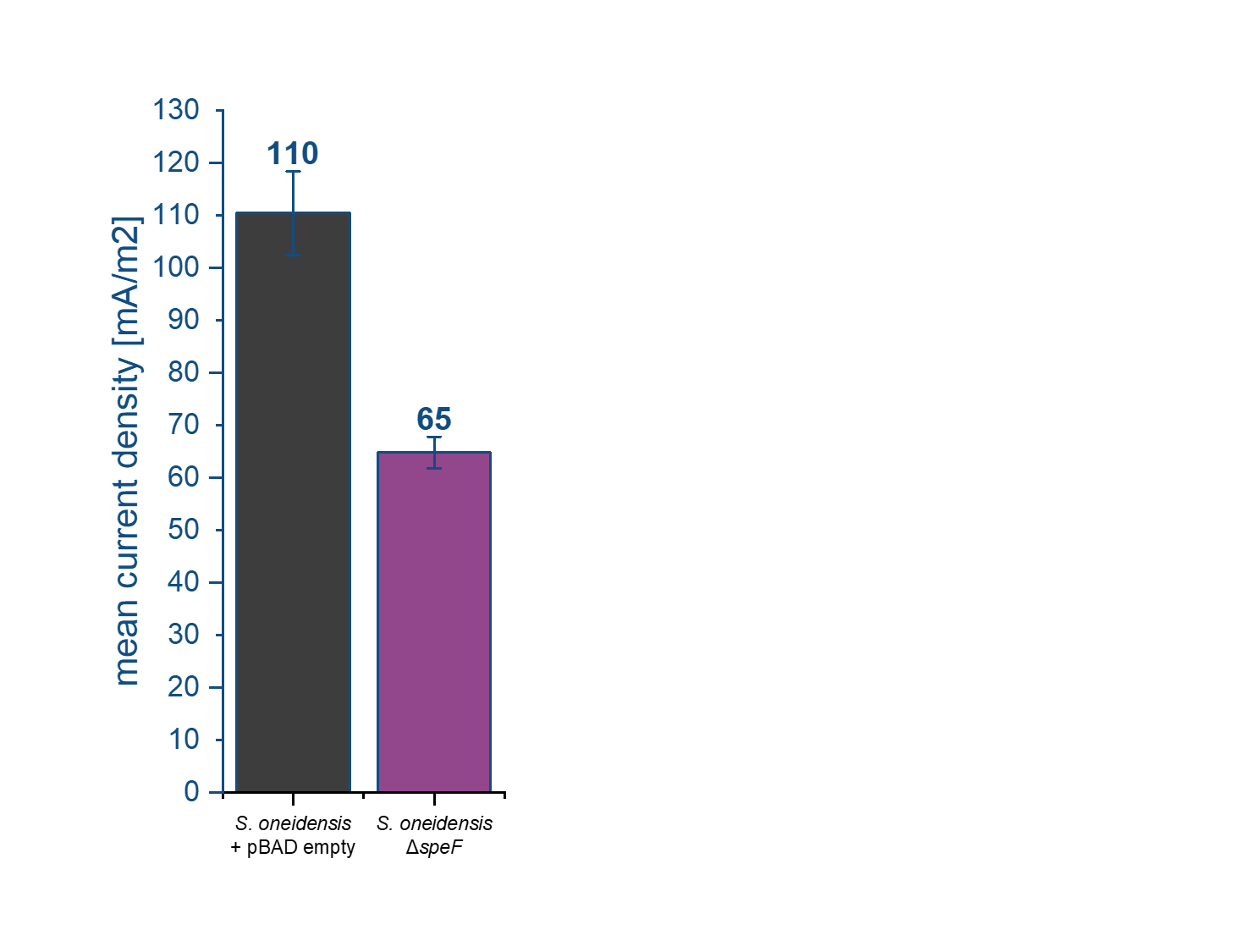

### Supplemental Figure 3

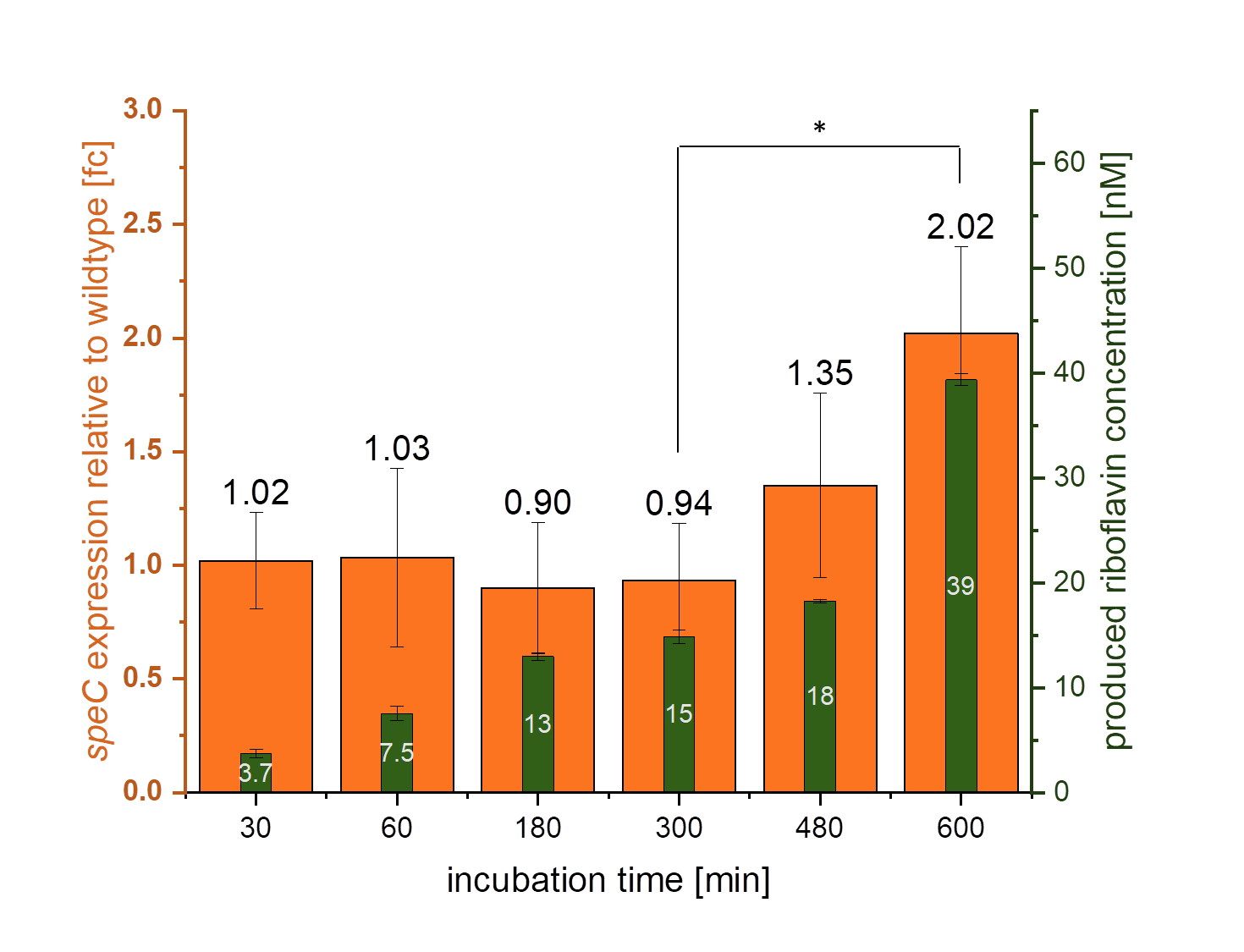

### Supplemental Figure 4

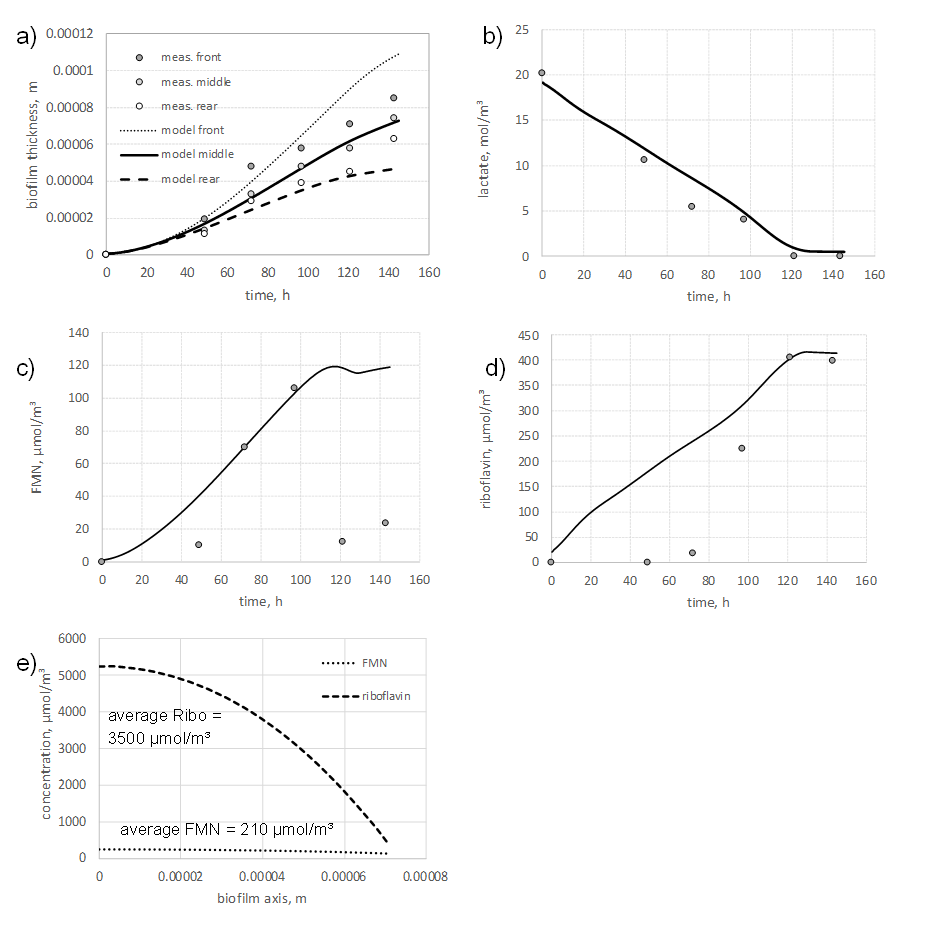
