## Supplemental Text for "Extracellular riboflavin induces anaerobic biofilm formation in *Shewanella oneidensis*"

### Supplements

Batch and biofilm model

Based on the experimental results from batch and continuous biofilm cultivation a model was formulated to identify the driving factors for riboflavin production both in batch and biofilm systems. As can be seen from figure 4 the production of riboflavin is directly linked to growth of *S. oneidensis*. A Monod based double saturation approach (lactate and fumarate) was used to describe cell growth and FMN production. The transformation of FMN to riboflavin is then described with a first order reaction kinetic again coupled to the two Monod terms for lactate and fumarate.

$\frac{\mathrm{dN}}{\mathrm{dt}}= \mu_{max} \frac{c_{Lac}}{{K_{S}+ c}_{Lac}} \frac{c_{Fum}}{{K_{S}+ c}_{Fum}} N \left[ \frac{cells}{m^{3}h} \right]$	(S1)

$\frac{dc_{Lac}}{\mathrm{dt}}= {- \mu}_{max}\frac{1}{Y_{cell/Lac}} \frac{c_{Lac}}{{K_{S}+ c}_{Lac}} \frac{c_{Fum}}{{K_{S}+ c}_{Fum}} N \left[ \frac{mol}{m^{3}h} \right]$	(S2)

$\frac{dc_{Fum}}{\mathrm{dt}}= {- \mu}_{max}\frac{1.3}{Y_{cell/Lac}} \frac{c_{Lac}}{{K_{S}+ c}_{Lac}} \frac{c_{Fum}}{{K_{S}+ c}_{Fum}} N \left[ \frac{mol}{m^{3}h} \right]$	(S3)

$\frac{dP_{FMN}}{\mathrm{dt}}= \mu_{max} Y_{FMN/cell} \frac{c_{Lac}}{{K_{S}+ c}_{Lac}} \frac{c_{Fum}}{{K_{S}+ c}_{Fum}} N \left[ \frac{\mu mol}{m^{3}h} \right]$	(S4)

$\frac{dP_{Ribo}}{\mathrm{dt}}= k_{Ribo} P_{FMN} \frac{c_{Lac}}{{K_{S}+ c}_{Lac}} \frac{c_{Fum}}{{K_{S}+ c}_{Fum}} \left[ \frac{\mu mol}{m^{3}h} \right]$	(S5)

*N* is the number of cells, *µ_max_* the maximum growth at 30 °C, *c_Lac_* the lactate concentration and *c_Fum_* is the fumarate concentration (equation S1). *Y_cell/Lac_* is the yield coefficient for produced cells per transformed lactate (equations S2 and S3). *P_FMN_* is the intermediate product (FMN) and *Y_FMN/cell_* is the growth related yield coefficient (equation S4). The latter is a Luedeking-Piret approach with growth related production of FMN (1). S5 is the first order reaction rate coupled to substrate availability. *k_Ribo_* is the respective reaction rate constant.

In table S3 the kinetic and stoichiometric parameters are shown both for the batch culture and the biofilm simulation.

Knowing the production of FMN and riboflavin in a batch culture the next step was the simulation of its production in a biofilm. A 1D model was used to describe both biofilm and product formation (2, 3). The model is based on diffusion coupled with reaction inside the biofilm. The simulation tool AQUASIM was used to solve the equations (Reichert, 1998). The equations for substrate utilization and product formation inside the biofilm are then

$\frac{\partial c_{Lac}}{\partial t} ={- D_{Lac} \frac{\partial^{2}c_{Lac}}{\partial z^{2}}- \mu}_{max}\frac{1}{Y_{cell/Lac}} \frac{c_{Lac}}{{K_{S}+ c}_{Lac}} N \left[ \frac{mol}{m^{3}h} \right]$	(S6)

$\frac{\partial P_{FMN}}{\partial t} ={- D_{FMN} \frac{\partial^{2}P_{FMN}}{\partial z^{2}}+ \mu}_{max} Y_{FMN/cell} \frac{c_{Lac}}{{K_{S}+ c}_{Lac}} N$	$\left[ \frac{\mu mol}{m^{3}h} \right]$	(S7)

$\frac{\partial P_{Ribo}}{\partial t} ={- D_{Ribo} \frac{\partial^{2}P_{Ribo}}{\partial z^{2}}+ k}_{Ribo} P_{FMN} \frac{c_{Lac}}{{K_{S}+ c}_{Lac}}$	$\left[ \frac{\mu mol}{m^{3}h} \right]$	(S8)

*z* is the vertical axis within the biofilm and *D* the diffusion coefficient, which can be calculated with the Wilke-Chang equation (4). However, the most sensitive parameters for the simulation of biofilm dynamics is, beside growth rate *µ_max_* and yield coefficients (here *Y_cell/lac_* and *Y_FMN/cell_*), the cell density within the biofilm (5). The latter is responsible on how quick the thickness of the biofilm will develop, or how much new grown cells do fit into a certain biofilm volume (5). In our case the density has been measured and could be introduced into the model (Fig. S4, Tab. S3). To describe the gradient along the flow cell used for biofilm cultivation, we divided available reactor volume and biofilm surface in three connected compartments (front, middle and rear).

It has already been shown that the release of product from aggregated cells is not always based on a simple diffusion process as formulated in equations S4 and S5. Partly products are fixed to the outer cell wall and can be released by pH modification Coles & Gross 1967 (6). The diffusion coefficient has been used as tool to fit the measured data of product release with a simulation (7). Here the diffusion coefficient *D_Ribo_* used for riboflavin (see Tab. S3) is three orders of magnitude smaller compared to the value calculated with the Wilke-Chang equation.

The simulation results in figure S4 show a good fit for biofilm thickness *L_F_* along the flow cell. Due to the small diameter of the flow cell and respective shear forces acting on the biofilm, detachment was assumed to depend on a maximum achievable biofilm thickness *L_F_max_* (150 µm):

$\frac{dL_{F}}{dt} = u_{F}*\frac{L_{F}}{L_{F max}}$	(S9)

*u_F_* is the velocity by which the biofilm surface is moving perpendicular to the substratum (2). The lactate removal has been simulated by mimicking the reduced availability of an electron acceptor (oxygen) with an increasing *K_S_* value (Fig. S4 b).

Both FMN and riboflavin release are not properly displayed with the model approach (Fig. S4 c and d). Although the maximum concentration in the bulk can be reproduced, the combination of FMN production, its transformation to riboflavin and the adsorption/release process within the biofilm cannot sufficiently be simulated over time. A significant difference of the kinetic values chosen for batch and biofilm cultivation can be seen for *k_Ribo_*. Whereas the value for batch cultivation is 0.14 1/d the riboflavin production within the biofilm can only be displayed with a *k_Ribo_* 120 1/d. The main reason is the two times higher cell density (Tab. S3).

The high diffusion coefficient chosen for riboflavin helps to fit the final concentration values measured both for FMN and riboflavin within the biofilm (Fig. S4 e). However, the maximum concentrations might be misleading. Finally, the amount of riboflavin, which is fixed in the biofilm grown in the flow cell after 140 h is only **4.6 x 10^-3^ nmol**. On the other side the amount of riboflavin transported out of the system with the volumetric flow is about **24 nmol**.

Table S1: Strains used in this study.

Table S2: Primers used in this study.

Fig.S1: Impact of putrescine addition on current generation and biofilm formation on anode surfaces.

Fig. S2: Impact of *speF* deletion on current density. As can be seen the current density decreases significantly as a result of s*peF* deletion*.*

Fig. S3: Impact of produced riboflavin on *speC* expression during growth. After 600 min 39 nM riboflavin are produced. As can be seen the relative *speC* expression doubles after 600 min.

Table S3: Parameters for simulation of *S. oneidensis* growth and riboflavin production both in batch and as continuous biofilm cultivation

Fig. S4: Simulation results for a) biofilm thickness (front, middle and rear of the flow cell), b) lactate, c) FMN and d) riboflavin concentration. e) Simulated distribution of FMN and riboflavin over the average biofilm thickness after 140 h.
