## Supplemental Table 1 for "Extracellular riboflavin induces anaerobic biofilm formation in *Shewanella oneidensis*"

Table S1: Strains used in this study.

| Strain | Source |
| --- | --- |
| *S. oneidensis* wildtype + pBAD empty | This study |
| *S. oneidensis* Δ*speC* + pBAD empty | This study |
| *S. oneidensis* wildtype + pBAD *speC* | This study |
| *S. oneidensis* Δ*speC* + pBAD *speC* | This study |
| *S. oneidensis* Δ *srtA* + pBAD empty | This study |
| *E. coli* WM3064 | W. Metcalf, Univ. of Illinois, Urbana |
| *E. coli* WM3064 +  pMQ500up_500down_*speC* | This study |
| *E. coli* WM3064 + pMQ500up_500down_*speF* | This study |
| *S. oneidensis* Δ*speF* | This study |
