## Supplemental Table 2 for "Extracellular riboflavin induces anaerobic biofilm formation in *Shewanella oneidensis*"

Table S2: Primers used in this study.

| **Num.** | **Name** | **Sequence** | **purpose** |
| --- | --- | --- | --- |
| 1 | *speC_*OL_pBAD_for | GTTTAACTTTAAGAAGGAGATATACATACCATGAGCCAATTTCAATCTATTG | Construction of pBAD *speC* |
| 2 | *speC_*OL_pBAD_rev | CCGCCAAAACAGCCAAGCTGGAGACCGTTTTTACAGATAGTAATCTTTCAGT | Construction of pBAD *speC* |
| 3 | pBAD_test_for | GATTAGCGGATCCTACCTGAC | Sequencing of pBAD inserts *speC* and *srtA* |
| 4 | pBAD_test_rev | CTCTCATCCGCCAAAACAGC | Sequencing of pBAD inserts *speC* and *srtA* |
| 5 | 500up_*speC*_OL_pMQ150_for | GATTACGAATTCGAGCTCGGTACCCGGGGATCTACTTCGACTCCAGTGAAGC | Construction of pMQ150 500up 500 down *speC* |
| 6 | 500up_*speC*_OL_500down_*speC*_rev | CTTTTTGTTGCCATTAAGCCAATGGGCCAGAGTCTCCTTTG | Construction of pMQ150 500up 500 down *speC* |
| 7 | 500down_*speC*_for | TTGGCTTAATGGCAACAAAAAG | Construction of pMQ150 500up 500 down *speC* |
| 8 | 500down_*speC*_OL_ pMQ150_rev | CGGCCAGTGCCAAGCTTGCATGCCTGCAGGCGTTAACAGACCCAATGCGT | Construction of pMQ150 500up 500 down *speC* |
| 9 | 500up_*srtA*_OL_pMQ150_ for | CAGTGCCAAGCTTGCATGCCTGCAGGGCGAGCCTAAGGAGTTAGT | Construction of pMQ150 500up 500 down *srtA* |
| 10 | 500up_*srtA*_rev | CGAGCTCATCAATGGCTAAG | Construction of pMQ150 500up 500 down *srtA* |
| 11 | 500down_*srtA*_OL_500up_*srtA_*for | CATTGATGAGCTCGACGGACGATAACGCGGCAGA | Construction of pMQ150 500up 500 down *srtA* |
| 12 | 500down_*srtA*_OL_ pMQ150_rev | TTACGAATTCGAGCTCGGTACCCGGGGATCGACCGACGTAAAGACCAAC | Construction of pMQ150 500up 500 down *srtA* |
| 13 | 500up_prpR_for_OL_pMQ | CGAATTCGAGCTCGGTACCCGGGGATCACAACTAACGGATAGCGCCA | Construction of pMQ150 500up 500 down *prpR* |
| 14 | 500up_prpR_rev_OL_500d_prpR | CGATGCTTATGGGCTGCGACGCTGCGCATCACTTATTGATTGC | Construction of pMQ150 500up 500 down *prpR* |
| 15 | 500down_prpR_for | GTCGCAGCCCATAAGCATCG | Construction of pMQ150 500up 500 down *prpR* |
| 16 | 500down_prpR_rev_OL_pMQ | CGGCCAGTGCCAAGCTTGCATGCCTGCAGGCTTCAGTGCTAACAACGGCT | Construction of pMQ150 500up 500 down *prpR* |
| 17 | 500up_prpB_for_OL_pMQ | CGAATTCGAGCTCGGTACCCGGGGATCCTCAACTCACAGTGAGTGAG | Construction of pMQ150 500up 500 down *prpB* |
| 18 | 500up_prpB_rev_OL_500d_prpB | CCATGGTCTGTACTTATCTTGGTCTTGTCCTTATCCAATAAAAAG | Construction of pMQ150 500up 500 down *prpB* |
| 19 | 500down_prpB_for | AAGATAAGTACAGACCATGGACG | Construction of pMQ150 500up 500 down *prpB* |
| 20 | 500down_prpB_rev_OL_pMQ | CGGCCAGTGCCAAGCTTGCATGCCTGCAGGCGCTCTAACACTTCTTTTAATG | Construction of pMQ150 500up 500 down *prpB* |
| 21 | Test_Δ*speC*_for | GCCTAAGCGTAATGCCGA | Sequencing of Δ*speC* |
| 22 | Test_Δ*speC*_rev | TGCAAGAATTACCCTTTGCG | Sequencing of Δ*speC* |
| 23 | Test_Δ*srtA*_for | CGGTGCTGACCGATCTGCA | Sequencing of Δ*srtA* |
| 24 | Test_Δ*srtA*_rev | ACCAACTGGGCGATCAAGT | Sequencing of Δ*srtA* |
| 25 | Test_Δ*prpR/prpB*_for | CAGTACCTCAGGTCGATGCC | Sequencing of Δ*prpR* and Δ*prpB* |
| 26 | Test_Δ*prpR/prpB*_rev | CGGCAATTTGGCTTTGCTCGC | Sequencing of Δ*prpR* and Δ*prpB* |
| 27 | qPCR*_S.on*_for | TATTCAAGTGCTTCTATTAG | Cell quantification of *S. oneidensis* via qPCR |
| 28 | qPCR_*S.on*_rev | AAGAACTTCTACTCAACA | Cell quantification of *S. oneidensis* via qPCR |
| 29 | qPCR_*S.on_speC_*for | GTGAAGAGCATCTTCGACC | *S. oneidensis speC* quantfication via qPCR |
| 30 | qPCR_*S.on*_*speC_*rev | TAGTTAGCAGGGAAGCCG | *S. oneidensis speC* quantfication via qPCR |
| 31 | qPCR_*S.on_rpoA*_for | GAAGCTATCCGTCGTTCTGC | *S. oneidensis rpoA* quantfication via qPCR |
| 32 | qPCR_*S.on_rpoA*_rev | GACAGGACGCAACAGAATCG | *S. oneidensis rpoA* quantfication via qPCR |
| 33 | 500up_*speF*_OL_pMQ150_for | GTCACGACGTTGTAAAACGACGGCCAGTGCCAAGCTTGCATGCCTGCAGGAAAGTTTGCGACTGTGGCTA | Construction of pMQ150 500up 500 down *speF* |
| 34 | 500up_*speF*_rev | TTGAACCTGACTGCAGAGTTA | Construction of pMQ150 500up 500 down *speF* |
| 35 | 500down_*speF*_OL_500up_*speF*_for | GAAATTGACTGCAGAGCGGGGTTATTAACTCTGCAGTCAGGTTCAACTAGTCGTTAATTTCCTTTTTACTTTAAAA | Construction of pMQ150 500up 500 down *speF* |

| 36 | 500down_*speF*_OL_pMQ150_rev | GGAAACAGCTATGACCATGATTACGAATTCGAGCTCGGTACCCGGGGATCATCATATTTTTATTGATAACACAC | Construction of pMQ150 500up 500 down *speF* |
| --- | --- | --- | --- |
| 37 | Test_Δ*speF*_for | CAACCGATCAATGACACGC | Sequencing of Δ*speF* |
| 38 | Test_Δ*speF*_rev | CAGTCAATCATCCTTAAGTATC | Sequencing of Δ*speF* |
