## Supplemental Table 3 for "Extracellular riboflavin induces anaerobic biofilm formation in *Shewanella oneidensis*"

Table S3: Parameters for simulation of *S. oneidensis* growth and riboflavin production both in batch and as continuous biofilm cultivation

| Parameter | Value batch  experiments | Values for cultivation as biofilm |
| --- | --- | --- |
| Growth rate *µ_max_* h^-1^ | 0.35 | 0.34 |
| Monod constant *K_S_* mol m^-3^  Yield coefficient *Y_cell/Lac_* cells mol^-1^ | 22  1.3 x 10^13^ | 25 – 120  7.0 x 10^11^ |
| Product yield *Y_FMN/cell_* µmol cell^-1^ | 9.0 x 10^-13^ | 3.9 x 10^-11^ |
| FMN 🡪 Ribo *k_Ribo_* h^-1^ | 0.14 | 120 |
| Cell density cells m^-3^ | 8.0 x 10^14^ | 5.15 x 10^16^ |
| Diffusion coefficient: |  |  |
| Riboflavin *D_Ribo_* m^2^ h^-1^  FMN *D_FMN_* m^2^ h^-1^ | -  - | 3.0 x 10^-9^  1.9 x 10^-6^ |
| Lactate *D_Lac_* m^2^ h^-1^ | - | 4.2 x 10^-6^ |
